# Evaluation and application of chemical decrosslinking in the context of histopathological spatial proteomics

**DOI:** 10.64898/2026.02.06.704439

**Authors:** Andikan J. Nwosu, Liang Chen, Rashmi Kumar, Yumi Kwon, Shaun M. Goodyear, Adel Kardosh, James M. Fulcher, Ljiljana Paša-Tolić

**Affiliations:** Environmental Molecular Sciences Laboratory, Pacific Northwest National Laboratory, Richland, Washington, United States; Knight Cancer Institute, Portland, Oregon, United States; Department of Medicine, Division of Medical Oncology, Oregon Health & Science University, Portland, Oregon, United States

## Abstract

Laser capture microdissection (LCM) - based spatial mass spectrometry proteomics is a rapidly emerging technique with strong potential for use in formalin-fixed, paraffin-embedded (FFPE) tissues. Several sample-preparation methods have been developed to decrosslink FFPE proteins for spatial proteomics; however, residual crosslinks often remain, and depth can remain impaired relative to fresh frozen tissue samples. To increase proteome coverage in spatially resolved LCM-FFPE samples, we investigated a panel of chemical compounds with the potential to catalyze the decrosslinking of nucleophilic functional groups on proteins. Systematic screening and optimization of temperature, incubation time, and reagent concentration led to the identification of 3,4-diaminobenzoic acid as an effective agent for improving proteome coverage in FFPE pancreatic tissue. This compound could boost precursor identifications by more than 10% at both reduced (70 °C) and high (90 °C) temperatures. Application of this chemical-decrosslinking strategy to a pancreatic ductal adenocarcinoma tissue section enabled the identification of numerous cell-type–enriched proteins with clinical and therapeutic relevance. Taken together, our findings show that chemical decrosslinking can increase proteome coverage in FFPE tissues, thereby advancing our understanding of tissue microenvironments in physiological and pathological contexts.

## INTRODUCTION

Formalin-fixed, paraffin-embedded (FFPE) tissues have long been a cost-effective preservative method for patient tissues in pathology archives. Apart from their use in histology or immunohistochemistry, FFPE tissues are beneficial to translational research.^1^ FFPE samples have been used in genomic, transcriptomic, and proteomic analyses due to their accessibility and (in many cases) the inclusion of associated pathology reports, diagnosis, treatments, and clinical outcomes. The fixation process also reduces sample degradation during handling and transportation from one facility to another.^2^ Mass spectrometry (MS)-based proteomic analysis of FFPE tissues plays a crucial role in evaluating medical treatments,^3^ discovering biomarkers,^4^ advancing drug development,^5^ and understanding disease mechanisms^6^ owing to the depth and sensitivity of MS proteomics in detecting subtle changes in protein abundance.

Spatial proteomics has increasingly used FFPE tissue to obtain a global view of intact tissue samples with cell-type or functional unit (i.e., pancreas islet) resolution. However, proteome coverage from FFPE tissues is often significantly lower relative to fresh frozen (FF) tissues due to the presence of intra- and inter-molecular protein crosslinks.^7,8^ Therefore, the efficient removal of aldehyde crosslinks from proteins is essential for protein retrieval, maximizing proteome depth, and ensuring accurate identification and quantification - particularly for low input, near single cell level samples.

The advancement of approaches that improve detection of proteins in FFPE tissues started decades ago in the context of immunohistochemistry (IHC) applications.^9^ The development of heat-induced antigen retrieval techniques specifically provided a path towards widescale adoption of protein measurements in FFPE tissue with IHC.^10^ The main discovery and rationale behind this technique was the use of high temperature for reversing crosslinks to expose masked functional groups in antibody ligands thereby enabling antibody binding on the tissues for improved staining efficiency.

This rationale has been employed in sample preparation for mass spectrometry bottom-up proteomics (BUP) as well. High temperature-based protein lysis has been used extensively in FFPE proteomics studies which have generally found improved protein identifications from such tissue samples.^11 - 13^ Furthermore, many clinical and translational studies involving FFPE tissues have benefited from the improved proteome coverage from high temperature sample preparation.^14 - 18^ As spatial proteomics has become more widely adopted for analyzing biological heterogeneity, the amount of cellular material being measured has correspondingly decreased with new low-input analytical approaches.

More specifically, techniques that employed laser-capture microdissection (LCM) have demonstrated the ability to achieve near single-cell BUP from tissue sections.^19 – 21^ Equally important has been the miniaturization of sample processing into the nanoliter regime, which reduces protein losses from nonspecific adsorption while improving digestion kinetics. For example, the nanodroplet processing in the one pot for trace samples (nanoPOTS) workflow was adapted for the proteomic analysis of LCM-isolated FFPE samples.^22^ In this method, protein decrosslinking and extraction were achieved by heating the samples to 90 °C for 90 minutes in the presence of an MS-compatible extraction buffer including 0.1% of *n*-dodecyl-β-d-maltoside (DDM). This approach yielded up to 88% proteome coverage from FFPE mouse liver tissue. This level of coverage is comparable to results typically achieved with FF tissue, with approximately 10–12% loss likely due to remaining cross-links.^22^

Inspired by the organocatalytic removal of formaldehyde adducts from RNA and DNA bases,^23^ we hypothesized that a mass spectrometry compatible compound could similarly be incorporated into the nanoPOTS sample preparation workflow to facilitate protein decrosslinking, thereby further improving proteome coverage for near single cell FFPE samples. Through a literature review and generalization of identified factors that contribute to decrosslinking,^23 – 28^ four chemical compounds were selected for assessment. This included Bismuth Tribromide (BiBr3) and 2-Amino-5-methylbenzoic acid which were identified from the literature, as well as 3,4-diaminobenzoic acid and *o*-phenylenediamine which we hypothesized to be functionally similar based on their chemical structure. Compounds that passed initial criteria (such as aqueous solubility) were introduced during protein extraction of 100 µm x 100 µm square regions of a 10µm thick FFPE pancreatic tissue section. Of the compounds evaluated, 3,4-diaminobenzoic acid was found to provide the highest number of precursor and protein identifications. A deeper examination revealed a striking increase in recovery of peptides containing cysteine residues, suggesting that the mechanism of increased decrosslinking is particularly effective at thiol crosslinks. Furthermore, adding 3,4-diaminobenzoic acid at reduced temperature (70°C) increased the protein identification by 24% (i.e., from 2,639 to 3,283 proteins). The use of reduced temperature conditions for decrosslinking FFPE tissues is particularly advantageous for nanoliter-scale applications where preventing excessive evaporation can be challenging. We further applied the full decrosslinking workflow to study several cell types isolated from a pancreatic ductal adenocarcinoma (PDAC) tissue section. We demonstrated enrichment of cancer and non-cancerous protein markers across 5,700 unique proteins tested for differential abundance. Taken together, our work described how the introduction of 3,4-diaminobenzoic acid during protein extraction from FFPE tissues can increase the proteome coverage thereby expanding the potential for greater discoveries in clinical proteomics applications.

## EXPERIMENTAL SECTION

### Reagents and materials

Dithiothreitol (DTT), 2-chloroacetamide (CAA), and 2,2,4-tri-methylpentane were obtained from Thermo Fisher Scientific (Waltham, USA). 3,4-diaminobenzoic acid (Compound A) was purchased from TCI (Tokyo, Japan). n-dodecyl-β-D-maltoside (DDM), ammonium bicarbonate (ABC), anhydrous acetonitrile (ACN), ethanol (EtOH), dimethyl sulfoxide (DMSO), formic acid (FA), 2-Amino-5-methylbenzoic acid (Compound B), *o*-Phenylenediamine (Compound C), and heptadecafluoro-1,1,2,2-tetrahydrodecyldimethylchlorosilane (PFDS) were obtained from Sigma-Aldrich (St. Louis, USA). Trypsin (MS grade) and Lys-C (MS grade) were purchased from Promega (Madison, USA). Deionized water (18.2 MΩ) was produced using a Barnstead Nanopure Infinity system (Los Angeles, USA).

### FFPE Tissue Sections

Normal pancreatic FFPE tissue block was purchased from US Biolab Inc. (Rockville, MD). The PDAC tumor was obtained from the Breden-Colsen Center for Pancreatic Care at Oregon Health and Science University (OHSU), United States. This study was approved by OHSU Institutional Review Board (IRB#28644). The tissue was sectioned into 10 μm slices using a Leica microtome and then placed onto PEN membrane slides. The FFPE tissue slides were stored at 4 °C until further use. To dewax, the tissue slides were immersed in xylene twice for 1 minute each, followed by rehydration through a series of ethanol solutions: 100% ethanol for 15 seconds, 95% ethanol for 15 seconds, 70% ethanol for 15 seconds, and deionized water for 10 seconds. After rehydration, the tissues sections were stained with hematoxylin for 5 seconds, rinsed in deionized water for 10 seconds, and then dehydrated in 70% ethanol for 10 seconds. The sections were counterstained with eosin through a quick dip, followed by dehydration in 95% ethanol for 15 seconds, 100% ethanol for 15 seconds, and xylene for 10 seconds. After processing and staining, the FFPE slides were dried in a vacuum desiccator and stored at 4 °C for further use.

### Laser Capture Microdissection (LCM)

Prior to the LCM experiments, 200 nL of DMSO was preloaded into each nanowell of the nanoPOTS chip to serve as a capture solvent for tissue collection, as described in our previous study.^29^ A PALM MicroBeam system (CarlZeiss MicroImaging, Munich, Germany) was utilized for collecting samples from FFPE tissues slides using the ‘CenterRoboLPC’ function. The nanoPOTS chip was mounted onto the LCM system using a slide adapter (SlideCollector 48, Carl Zeiss MicroImaging). Square tissue regions of interest (ROI) with a side length of 100 μm were selected and collected at 5 × magnification, with a cutting energy of 48, and a focus setting of 88. To capture the isolated tissue into the capture droplet, laser pressure catapulting (LPC) energy of 20 and focus of 5 was used. The success of the sample collection was confirmed by observing the tissue voxels within the DMSO droplet under the microscope. After sample collection, the nanoPOTS chip was incubated at 70°C to remove the capture droplets and stored at −20°C for further sample preparation.

### Proteomic sample preparation

A custom-built nanoliter-scale robotic liquid-handling platform was used for high throughput sample preparation, with some modifications to our previously established protocol.^30,31^ For sample preparation conditions, 200 nL of extraction buffer (1 mM DTT, 0.1% DDM, 100 mM ABC buffer, and 10% DMSO) was added to each nanowell, followed by incubation at 90 °C for 90 min to facilitate protein decrosslinking, extraction, and denaturation for standard high temperature condition. For other sample preparation with the compounds, the ABC buffer concentration was adjusted to ensure the decrosslinking buffer maintained a pH of 8.0 with a 5mM concentration of listed compounds in the protein lysis mix. This concentration was selected based on the work of Karmakar et al., 2015 with compound B.^23^ Therefore, 200 nL of extraction buffer with 5mM of 3,4-diaminobenzoic acid (referred to as Compound A), 2-Amino-5-methyl-benzoic acid (Compound B), or *o*-phenylenediamine (Compound C), 1mM DTT, 0.1% DDM, 100mM ABC buffer and 10% DMSO were added to each nanowell, followed by incubation at 90 °C for 90 min to facilitate protein decrosslinking, extraction, and denaturation for each compound test respectively. Next, the nanoPOTS chip was incubated at 70 °C for 15 min to evaporate the DMSO. Then, 100 nL of alkylation buffer (10 mM CAA in 50 mM ABC buffer) was added and incubated in the dark for 30 min at room temperature. Subsequently, 50 nL of digestion buffer (containing 0.01 ng nL^-1^ Lys-C and 0.04 ng nL^-1^ trypsin in 50 mM ABC buffer) was added and incubated at 37 °C for 10 h. After digestion, 50 nL of 5% FA was added and incubated at room temperature for 30 min to terminate the enzymatic reaction. The samples were dried in a desiccator and stored at – 20 °C for LC-MS/MS analysis.

### nanoLC-MS/MS analysis

Samples on nanoPOTS chips were injected directly from nanowells to an in-house-built nanoLC-MS/MS system.^31^ Peptides were first loaded onto a solid phase extraction (SPE) column (100 µm i.d., 4 cm length) packed with 5 µM, C18 packing material (300 Å pore size; Phenomenex) for online cleanup using mobile phase A (0.1% formic acid in water) for 5 minutes. Subsequently, the cleaned peptides were reverse washed and transferred to an analytical column (50 µm i.d., 25 cm length), which was in-house packed with 1.7 µm C18 material (BEH 130 Å pore size, Waters) for separation. The gradient began at 2% B, increasing to 8% B within 1 min, followed by a linear rise to 22% B over the next 29 min and a further increase to 35% B within 9 min. The gradient was then raised to 80% B in 1 min and held for 8 min before being reduced to 2% B in 1 min. It was then gradually increased to 35% B over 5 min. Finally, the column was flushed with 2% B for 5 min to re-equilibrate the system.

timsTOF SCP mass spectrometer (Bruker Daltonics, Bremen, Germany) was used to collect the data operating in DIA-PASEF mode. The separated peptides were ionized by a CaptiveSpray ion source with the capillary voltage set to 1600V, and dry gas at 3.0L/min with a dry temperature of 200 °C.

Peptides were analyzed with high-sensitivity mode enabled and the singly charged precursor ions were filtered with a polygon filter. Full MS1 data were acquired in a range of 100 – 1700 m/z and 1/K0 = 0.6Vs cm^-2^ to 1.6Vs cm^-2^. DIA windows range from 300 m/z to 1000 m/z with 25Th isolation windows and were acquired with ramp times of 166 ms. The collision energy ranged from 59 eV at 1/K0 = 1.6Vs cm^-2^ to 20 eV at 1/K0 = 0.6 Vs cm^-2^.

### Data Analysis

Raw files were processed with DIA-NN^33^ version 2.2.0 (academic license) and searched against the *Homo sapiens* proteome database downloaded from UniProtKB (accessed 09/12/24), including common contaminants such as keratins and albumins. *In silico* library prediction settings included fully tryptic peptides with 2 missed cleavages, peptide length 7 – 30 amino acids, precursor range from 350 – 1000 m/z, precursor charge 2 – 4, and fragment ion range of 200 – 1800 m/z. Static modification included cysteine carbamidomethylation (except for PDAC samples) and N-terminal methionine excision, as well as the variable modifications: oxidation of methionine (UniMod: 35), N-term acetylation (UniMod: 1), formaldehyde adducts on tyrosine or tryptophan (UniMod: 1009), and formylation of lysine (UniMod: 122). Maximum number of variable modifications was set to 2. Match between runs was enabled, normalization was disabled (enabled for PDAC samples), and MS1 and MS2 mass accuracy were set to 15 ppm. Other parameters not mentioned were left as default in DIA-NN software. The result files contained in pg.matrix and pr.matrix were exported into Perseus^34^ or imported into R prior to removal of identified contaminants. Precursor and protein identifications were median normalized prior to plotting Pearson correlations and coefficients of variation (CVs). Other figures were made within Microsoft Excel. Mass spectrometry data has been deposited to the ProteomeXchange Consortium via MassIVE repository with dataset identifier MSV000100017.

## RESULTS

### Comparison of decrosslinking compounds

We first tested a high temperature (90°C) and a reduced temperature (70°C) condition to determine temperature-based improvements in decrosslinking with FFPE pancreatic tissue. In line with prior results,^14 – 18^ our initial tests demonstrated a 32% increase in precursor identifications with higher temperatures (**Fig S1a** and **S1b**, *p*<0.0001). Therefore, all subsequent optimization experiments were carried out at 90°C as the baseline temperature. We next investigated compounds that might facilitate protein decrosslinking in FFPE tissues; specifically, 3,4-diaminobenzoic acid (Compound A), 2-amino-5-methylbenzoic acid (Compound B), *o*-Phenylenediamine (Compound C) and Bismuth Tribromide (BiBr3) (**Table 1**).

**Table 1.**
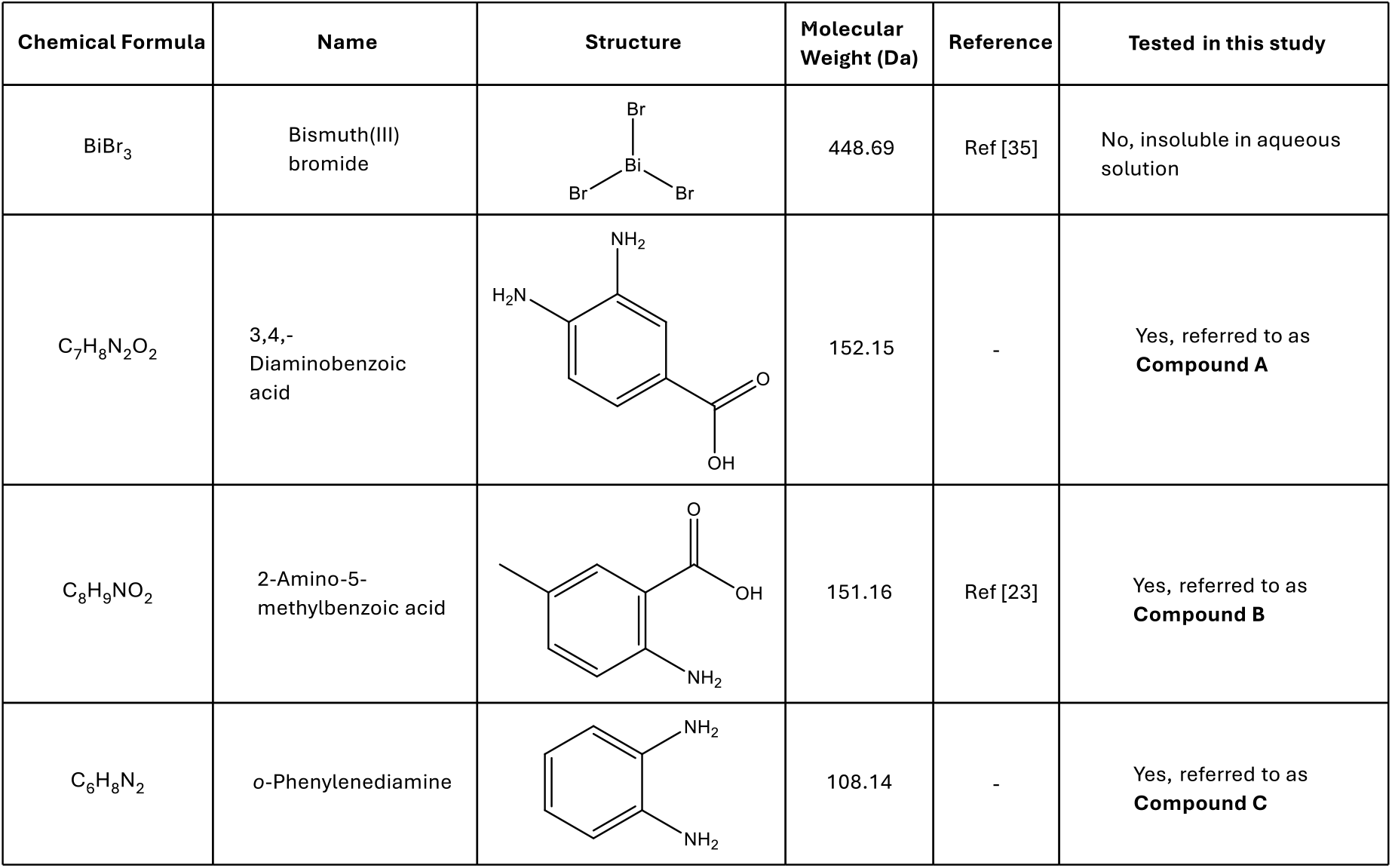
Table shows the list of compounds tested in this study; their physical properties and references used in screening these compounds.

| Chemical Formula | Name | Structure | Molecular Weight (Da) | Reference | Tested in this study |
| --- | --- | --- | --- | --- | --- |
| $\text{BiBr}_3$ | Bismuth(III) bromide | | 448.69 | Ref [35] | No, insoluble in aqueous solution |
| $\text{C}_7\text{H}_8\text{N}_2\text{O}_2$ | 3,4,-Diaminobenzoic acid | | 152.15 | - | Yes, referred to as **Compound A** |
| $\text{C}_8\text{H}_9\text{NO}_2$ | 2-Amino-5-methylbenzoic acid | | 151.16 | Ref [23] | Yes, referred to as **Compound B** |
| $\text{C}_6\text{H}_8\text{N}_2$ | o-Phenylenediamine | | 108.14 | - | Yes, referred to as **Compound C** |

Solubility of all compounds was first assessed at working concentrations of 5 mM, 0.5 mM and 0.1 mM. Unfortunately, BiBr3 was insoluble in water or PBS at each tested concentration and hence was not moved forward for testing (data not shown). Therefore, we focused on examining the three remaining compounds, the characteristics of which are described in Table 1. We prepared protein extraction buffer (0.1% DDM, 100mM ABC, 20% DMSO) with 5mM concentration for each of the three compounds and a fourth control condition without any compound in the mix (i.e., control). Sample preparation was carried out on 100 µm x 100 µm regions of a 10 µm thick pancreas FFPE tissue section using all four conditions. We observed a 14% and 13% increase in number of precursors using Compound A and Compound B respectively while there was a 17% decrease in number of precursors using Compound C compared to the standard high temperature sample preparation condition. This trend is also observed in protein identification with a 6% and 5% increase using Compound A and Compound B respectively and an 8% decrease with Compound C (**Fig 1a** and **1b**). An upset plot for protein similarities shows that on average 3,110 (80%) proteins and 19,044 (82%) precursors were identified in all four sample preparation conditions (**Fig 1d**). The precursor median coefficient of variation (CV) for four sample preparation conditions was 20%, 13%, 14%, 17% for standard high temperature, Compound A, Compound B, and Compound C respectively (**Fig 1c**), demonstrating that variation in these samples is at or below the expected norms in proteomics experiments.^36^

**Figure 1.**
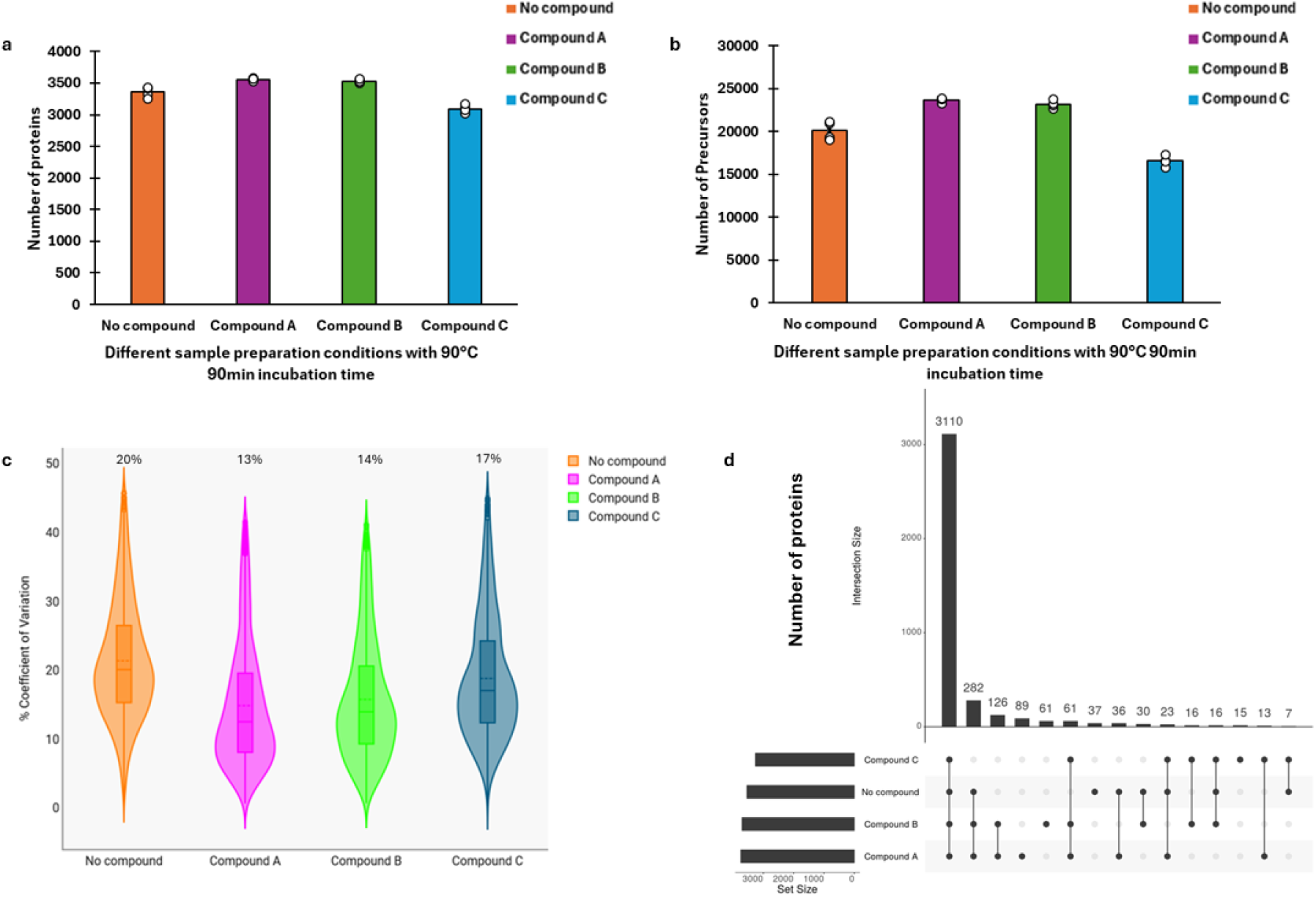
The number of proteins (**a**) and number of precursors (**b**) identified in all four sample prep conditions with compound A, B, and C in the protein lysis step in comparison to the standard high temperature prep. (**c**) Coefficients of variation (CV) of all four sample prep conditions. (**d**) Upset plot showing protein similarities between four sample prep conditions. N = 4 for each condition.

A Pearson correlation plot of protein intensities for all four sample preparation conditions show a correlation of 0.99 for each replicate analysis of standard high temperature (no Compound), Compound A, Compound B, and Compound C respectively (**Fig S2**). Lower correlation values could be due to heterogeneity across different sample collection regions. Gene ontology analysis using DAVID tool to classify the proteins similarities based on biological processes, revealed no significant differences in proteins identified (**Fig S3**). This indicates the different methods of sample preparation did not introduce bias toward any given process, function, or cellular component.

Crosslinks in FFPE tissue are primarily favored at nucleophilic amine groups (e.g., N-terminal α-amino and Lys ε-amines) as well as secondary crosslinks with arginine, glutamine, histidine, tryptophan, tyrosine, and cysteine sidechains.^37,38^ Consequentially, residual modifications often occur on tryptic peptides and previously crosslinked amino acid functional groups. Among these, prominent modifications include methylol adducts (Δ +27.99Da – note that formyl adducts may also be a product of exposure to formic acid in common LC-MS mobile phase) and a Schiff base or methylene bridge between two amino acids (Δ +12.00Da).^37,38^ Primary amine containing lysine residues appear to be particularly overrepresented with FFPE artifactual modifications.^40,41^ We sought to include a few of the most common modifications previously identified with DDA-database searches of FFPE datasets in order to see if they changed in abundance or frequency of identification in the presence of decrosslinking compound. In the Data Independent Acquisition (DIA) approach here, not every modification can reasonably be included as a variable modification during *in-silico* library generation due to the vast increase in library size and extensive search times associated with such databases. We also recognize that *in silico* predictions of these modifications may not be accurate due to underrepresentation in training datasets. Therefore, we restricted our search to a subset of modifications, specifically formyl modification of Lysine (K) due to this modification’s high-residue specificity and methylene/Schiff base modification of the low-frequency tryptophan (W) and tyrosine (Y) amino acids (**Fig 2a**). Common modifications such as methionine oxidation and N-terminal acetylation were included in the search as well. For formyl modification on Lysine precursor residues, we observed no significant differences in all four sample preparation conditions (Compounds A, B, C, and no compound). This is confirmed with *p* values of 0.87, 0.96, and 0.015 in Compound A, B, and C respectively (**Fig 2a**). Interestingly, cysteine-containing precursors were observed at significantly higher rates with compound aided sample preparations compared to standard high temperature sample preparation condition (**Fig 2b)**. Many of the cysteine-containing precursors were observed in proteins identified in the compound-containing samples, highlighting how some proteins can be quantified exclusively with compound treatment. (**Fig S4a** and **S4b**).

**Figure 2.**
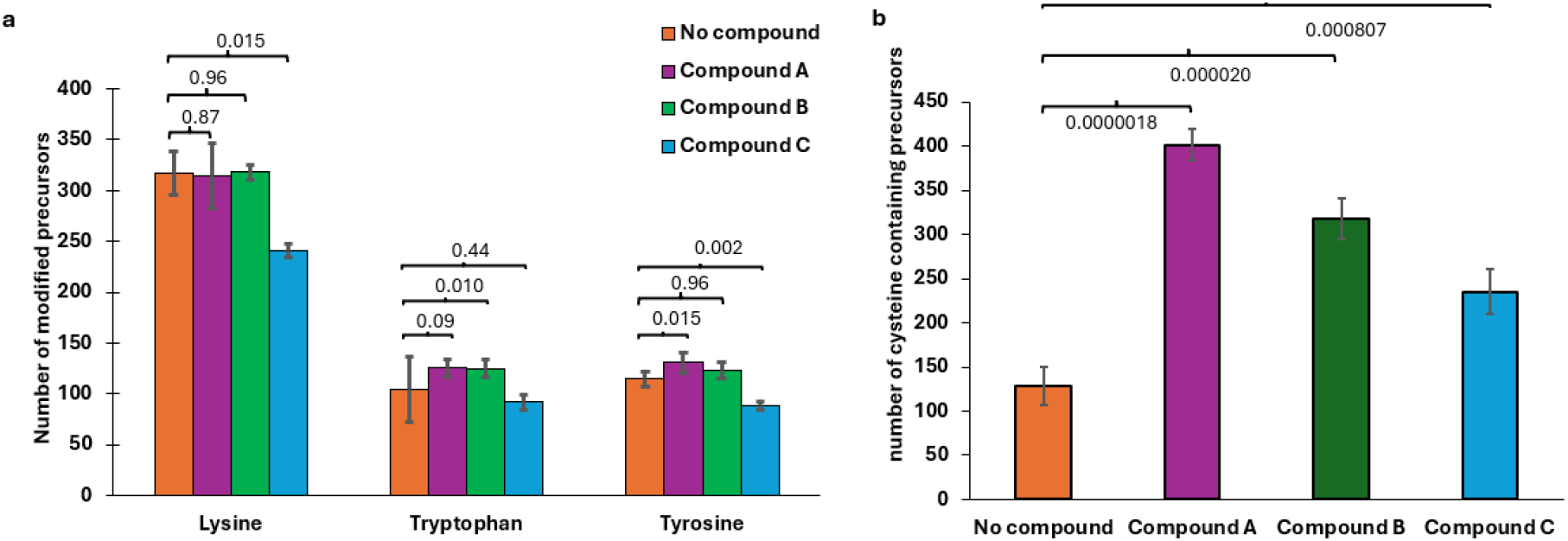
Precursor level amino acid residue modification (**a**) *p* values showing no significant differences in Lysine, Tryptophan, and Tyrosine modification in compound A, B, C compared to no compound. (**b**) *p* values showing significant differences in cysteine containing precursors in compound A, B, C compared to no compound.

These results suggest the decrosslinking compounds are particularly efficient at reversing a thiol-specific cysteine crosslink. Taken together, we suggest Compound A (3,4-diaminobenzoic acid) is the most efficient of the tested compounds for generally increasing precursor and protein identifications in FFPE samples.

### Optimization and general parameters of lead decrosslinking compound

Single cell and spatial proteomics require smaller sample processing volumes and surface area to reduce protein loss during sample preparation.^22^ In this study, sample processing volumes in the nanodroplet range coupled with high temperature can lead to increased evaporation. To cushion this effect, a humidifier was attached to the robotic platform.^22,31^ Given the difficulty in controlling evaporation in low volume samples, we desired to test the effect of adding Compound A at a lower temperature where evaporation would be minimized relative to 90°C. Our initial experiments with reduced temperature (70 °C) demonstrated corresponding reduction in proteome coverage (**Fig S1**). Considering the improved recovery of decrosslinked material using Compound A, we revisited the 70 °C incubation during the protein extraction and decrosslinking step. We observed a 19% increase in precursor identification with the use of Compound A (**Fig 3**). This demonstrates that Compound A can increase proteome coverage at both reduced and higher temperatures, making it amenable to experimental designs that utilize reduced incubation temperatures in order to limit nanodroplet evaporation.

**Figure 3.**
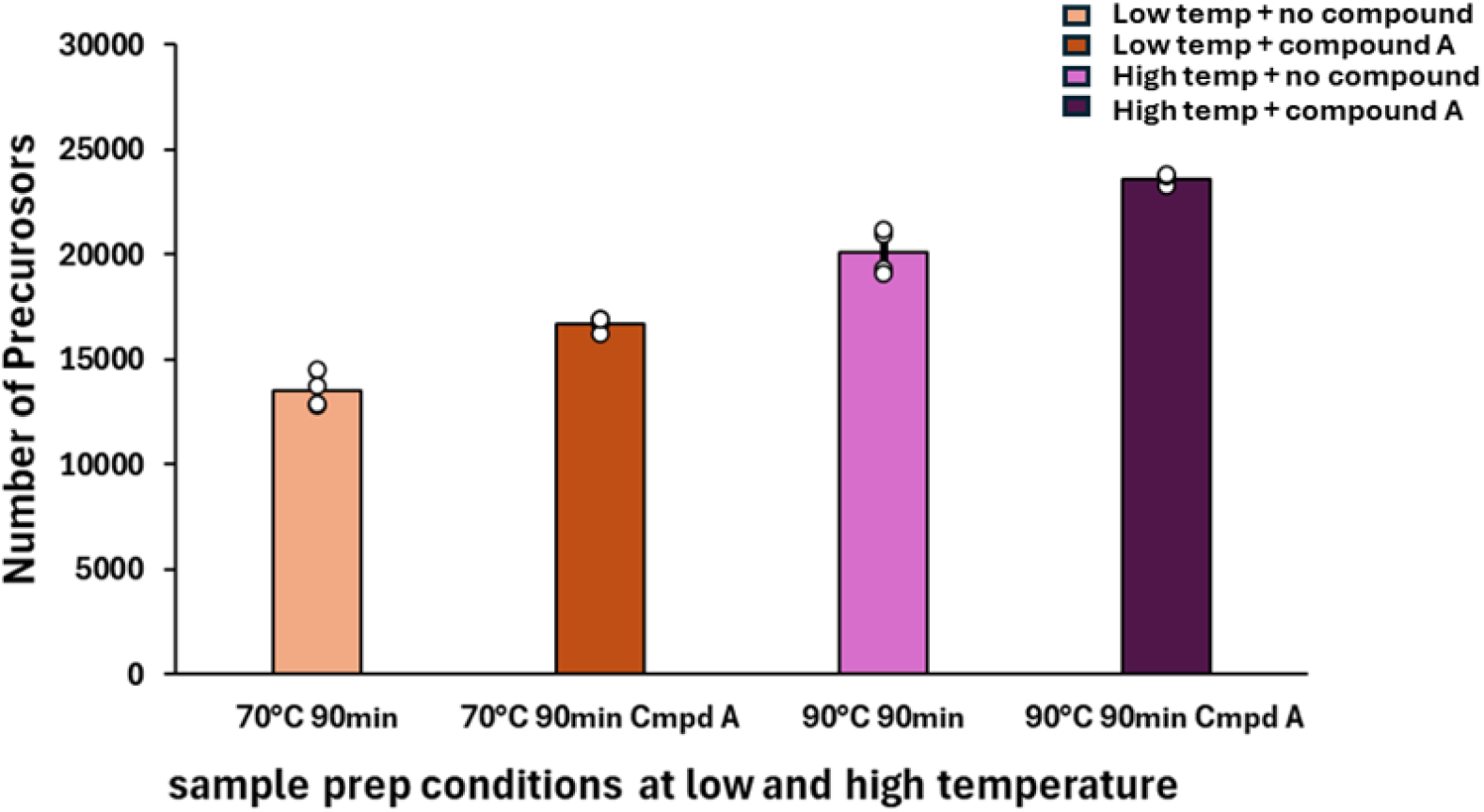
Effect of Compound A on overall precursor identifications, relative to control (N = 4), at 70°C and 90°C.

We further investigated the effect of concentration and time of incubation towards identifying the most favorable conditions at 90 °C. We prepared four different concentrations of compound A (1mM, 2.5mM, 5mM, 7.5mM) with 90 min incubation time and observed no significant difference in precursor and protein identification with the different compound A concentrations (**Fig S5a** and **S5b**). Given the catalytic effect of the compound, this data suggests substrate concentrations (i.e., protein crosslinks) are the limiting kinetic factor. We finally tested incubation time needed to obtain the highest protein yield. We prepared the samples and incubated at four different times (30, 60, 90, 120 mins) with 90 °C and 1mM concentration. We observed the highest protein yield at 120 min with an average of 28,772 precursors and 3,985 proteins identified. (**Fig S5c** and **S5d**).

### Application of full decrosslinking workflow to pancreatic ductal adenocarcinoma

Having identified a lead decrosslinking candidate compound and optimized parameters for low-input FFPE tissue samples, we next analyzed several cell types from a pancreatic ductal adenocarcinoma (PDAC) tissue section. Similar to the healthy human pancreas tissue above, we stained PDAC tissue with hematoxylin and eosin (H&E) to annotate different cell types. We utilized LCM to isolate ∼50,000 μm^2^ tissue areas corresponding to several cell types (**Fig 4**). This included pancreatic exocrine tissue (acinar, n = 3), ductal adenocarcinoma (cancer, n = 4), dysplastic ductal cells (dysplastic duct, n = 2), and stromal cells that were proximal (n = 3), or distal (n = 2) to tumor regions.

**Figure 4.**
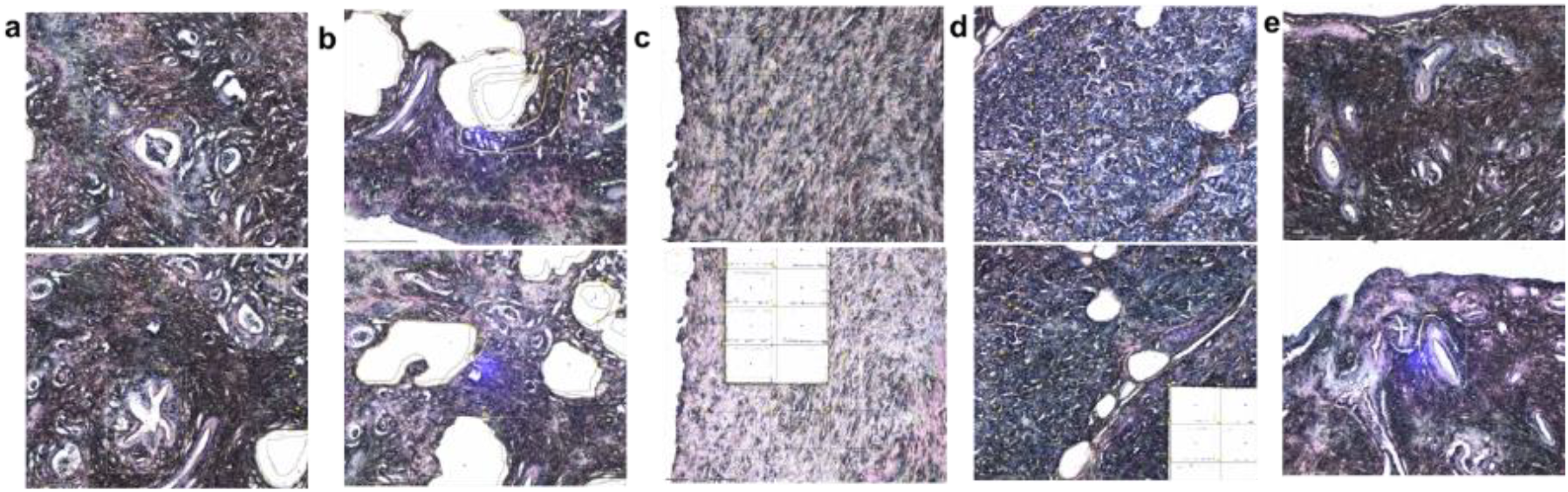
Representative images showing dissected regions from PDAC tissues for the following cell types: (a) cancer, (b) stroma proximal to cancer, (c) stroma distal to cancer, (d) acinar, and (e) dysplastic duct.

Applying the optimized FFPE tissue processing protocol (i.e., 120-minute incubation at 90 °C, 1 mM of Compound A) and nanoPOTS sample preparation, we identified 3,980 proteins on average across all samples analyzed with some variance depending on the cell type (**Fig 5a**). Encouragingly, this proteome coverage is close to the coverage obtained using healthy human pancreatic tissue, demonstrating the reproducibility of the method across different tissue types. Dimensional reduction using the 1,379 proteins with complete observations across the cell types could sufficiently resolve most cell types in principal component analysis, however the more closely related stromal cell types appeared to share feature variance from the first two principal components (**Fig 5b**). Differential abundance analysis provided several candidate proteins with enrichment for the various cell types (**Fig 5c**). Despite the relatively low statistical power from having few replicates, known proteins with cell type specificity could easily be identified and reached statistical significance (adjusted p.value <= 0.01). For example, PRSS1 (cationic trypsinogen), PRSS2 (anionic trypsinogen), CLPS (colipase), CPB1 (carboxypeptidase B), and CELA3A (chymotrypsin-like elastase family member 3A) are all enzymatic proteins known to be produced by acinar cells for executing exocrine pancreatic functions.^42,43^ Similarly for the pathological adenoma cell type, S100p (S100 calcium-binding protein P) and MYH14 (myosin heavy chain 14) have been shown to be enriched and play important roles in PDAC cell growth and aggressiveness.^44,45^

**Figure 5.**
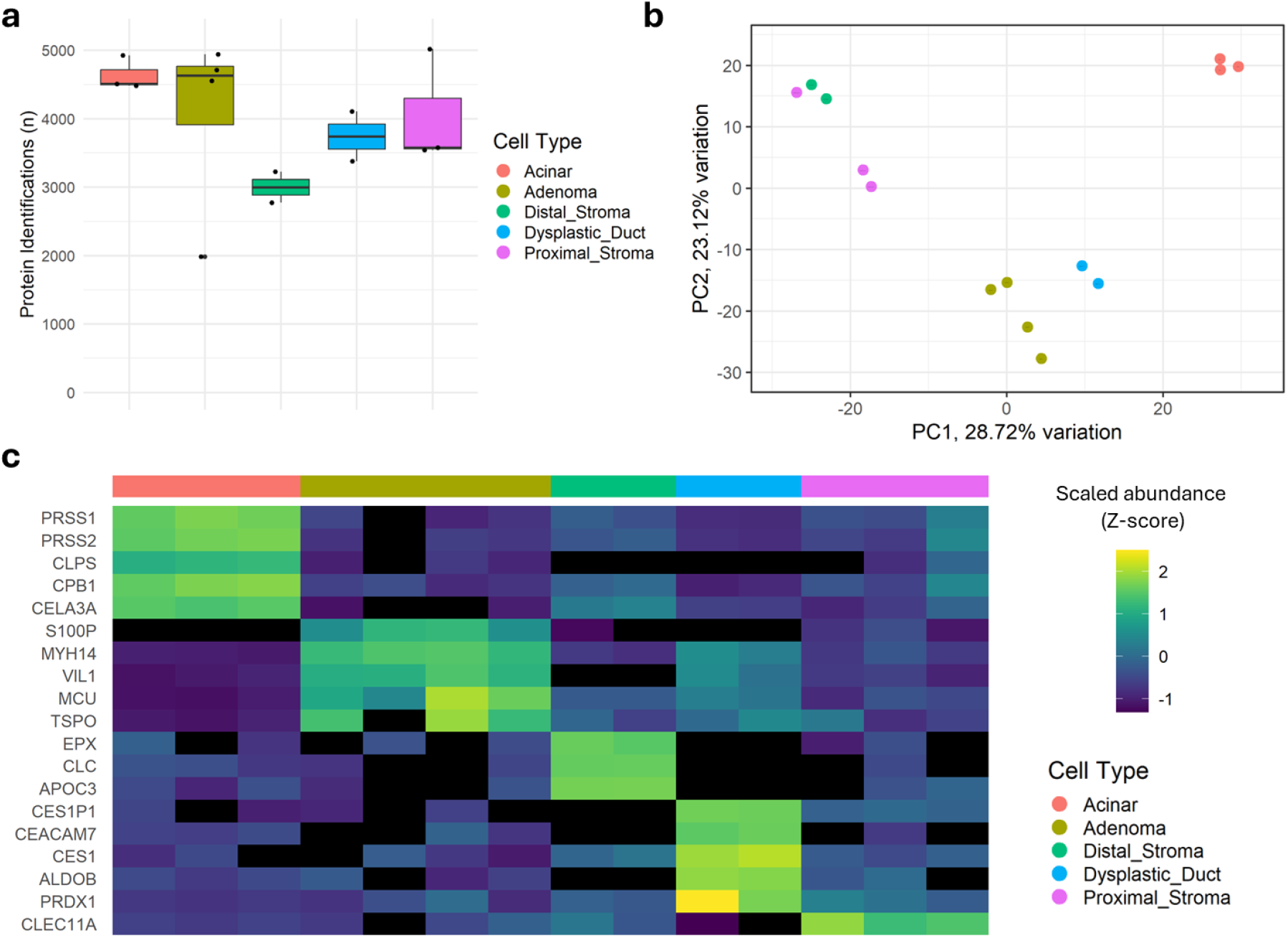
Application of decrosslinking workflow to pancreatic ductal adenocarcinoma (PDAC) tissue. (**a**) Boxplots showing the number of protein identifications for each cell type. (**b**) Principal component analysis of all PDAC samples, colored by cell type. Only proteins with complete observations (1,379 proteins) were used to generate dimensional reduction. (**c**) Heatmap of the top five, statistically significant cell type enriched proteins showing relative abundance across cell types. Statistically significant markers were determined using a two-sided, limma/Empirical Bayes t-test (adjusted p-values <= 0.01).

## DISCUSSION

FFPE tissues are an accessible and useful alternative to fresh frozen tissue samples for translational spatial proteomics. They represent a cost-effective means of storage for patient tissues, and large repositories exist in many hospitals and research facilities that can be used for research purposes. Several sample-preparation methods have been developed to decrosslink FFPE proteins for spatial proteomics; however, residual crosslinks often remain leading to impaired depth relative to fresh frozen tissue samples. Herein, we have demonstrated increased proteome coverage for low-input, LCM-FFPE spatial proteomics achieved with the aid of a chemical catalyst. We explored three chemical reagents and observed the highest proteome coverage with 3,4-diaminobenzoic acid. Generally, all compounds showed high reproducibility between replicates and no bias of proteome coverage across biological processes. Interestingly, cysteine-containing peptides appeared to be significantly increased in compound-treated samples, suggesting aldehyde-induced cysteine modifications are particularly susceptible to these compound’s decrosslinking effect. Furthermore, we optimized concentration, temperature, and incubation time to identify optimized conditions for spatial and low-input nanodroplet sample preparation methods. Specifically, the highest number of identifications could be achieved with 1mM 3,4-diaminobenzoic acid, 120 min incubation time, and 90 °C temperature. These optimized conditions were applied to a clinical PDAC tissue section towards identifying cell-type enriched proteins with oncological relevance. Using LCM, we isolated acinar, adenoma, stroma (proximal and distal to tumor), and dysplastic ductal cell populations prior to applying the optimized decrosslinking conditions. Although limited in sample size, many of the statistically significant proteins in acinar and adenoma cell types identified in this study have strong experimental support from prior studies.^34 - 37^ Of particular interest are dysplastic ductal cells which can be considered precursors to the adenoma cells in PDAC. Interestingly, we identified CEACAM7 as being enriched in dysplastic duct cells, and CEACAM7 has only recently been demonstrated as a potential therapeutic target in PDAC. CEACAM7 has been demonstrated to have high expression within the cancer stem cell-enriched subset of PDAC cultures, and CAR-T cells targeting CEACAM7 have shown efficacy against PDAC.^38^

## CONCLUSION

Chemical aided FFPE sample preparation reduces decrosslinks in FFPE tissues leading to increased proteome coverage. This is particularly advantageous with low-input FFPE samples, where near single-cell or single-cell resolved samples can have very low proteome coverage. Therefore, even modest improvements in proteome coverage can increase the potential of catching subtle proteomic changes associated with pathological states. Our study describes optimization of this chemical decrosslinking approach and demonstrates the potential in unraveling and identifying spatially resolved proteins with therapeutic or biomarker relevance.

## Acknowledgements

This research was supported through a Pacific northwest biomedical Innovation Co-laboratory (PMedIC) award to J.M.F., A.K., and L.P-T, and by the National Institutes of Health Grant UG3CA275697. A portion of the research was performed using EMSL (grid.436923.9), a DOE Office of Science User Facility sponsored by the Biological and Environmental Research program under Contract No. DE-AC05-76RL01830. We thank Drs. Eizabaru Sasatomi and Brian Mau for guiding identification of pathological areas in PDAC tissue for LCM collection; and the Knight BioLibrary and Histopathology Shared Resource for assistance in acquisition and sectioning of tissue sample. We thank Dr. Matthew Monroe for his assistance in depositing the raw proteomic data onto MassIVE.

## Author Contributions

**Andikan J. Nwosu** – Experimental design, sample preparation, data analysis and manuscript writing (initial draft).

**Liang Chen** – Experimental design, sample preparation, and manuscript writing (initial draft).

**Yumi Kwon** – Prepared and set up the LC-MS for data collection and collected data.

**Rashmi Kumar** – Experimental design, sample preparation, and formal analysis

**Shaun M. Goodyear** - Sample procurement, formal analysis, and manuscript review/editing

**Adel Kardosh** – Sample procurement, manuscript review/editing, and funding acquisition

**James M. Fulcher** – Manuscript writing, manuscript review/editing, funding acquisition, and formal analysis.

**Ljiljana Paša-Tolić**– Manuscript review/editing and funding acquisition. Final manuscript read and approved by all authors.

