## Supplementary Figures for "Evaluation and application of chemical decrosslinking in the context of histopathological spatial proteomics"

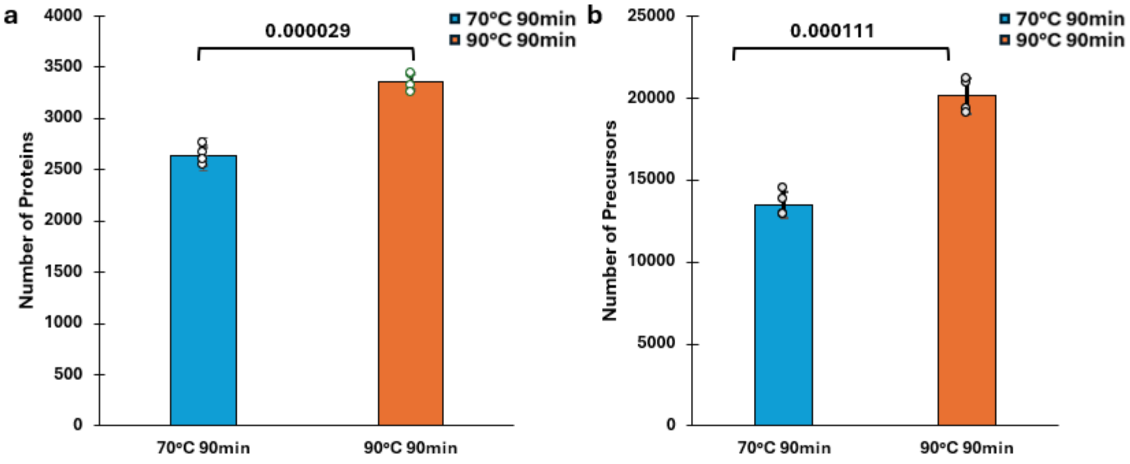


**Figure S1**. (**a**) Number of proteins and number of precursors (**b**) identified from 100 µm x 100 µm FFPE pancreas tissue sample (N = 4) at 70°C and 90°C.

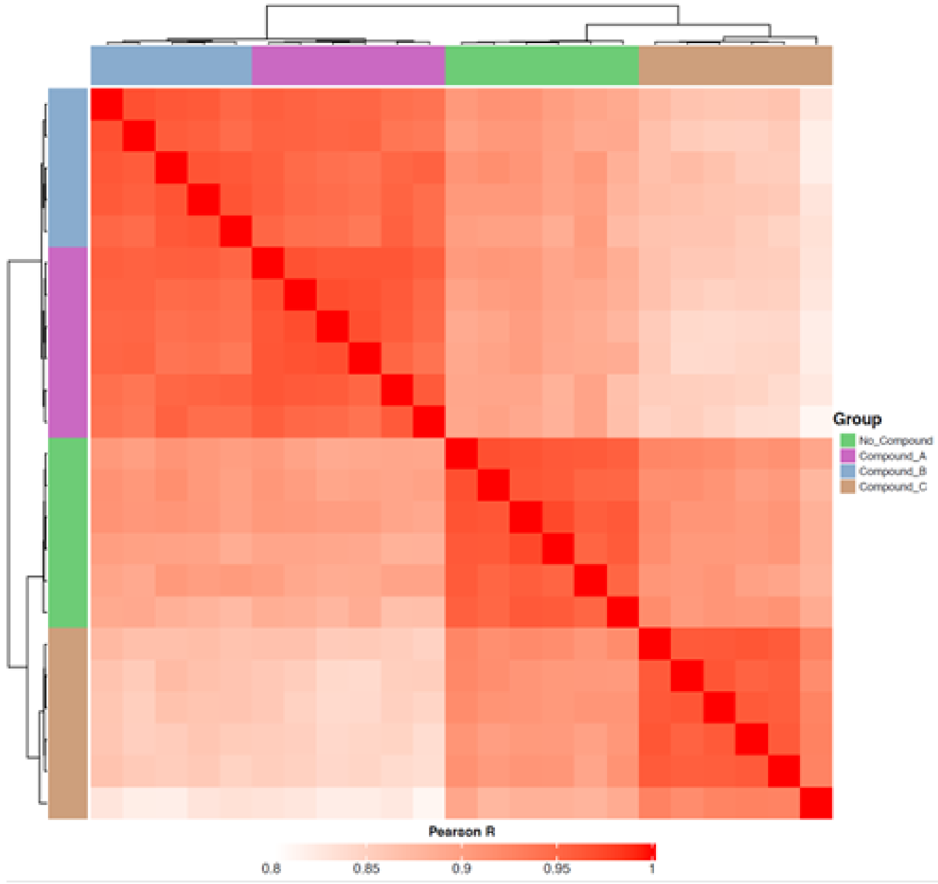


**Figure S2**. Pearson correlation of between samples and compounds tested (N = 4).

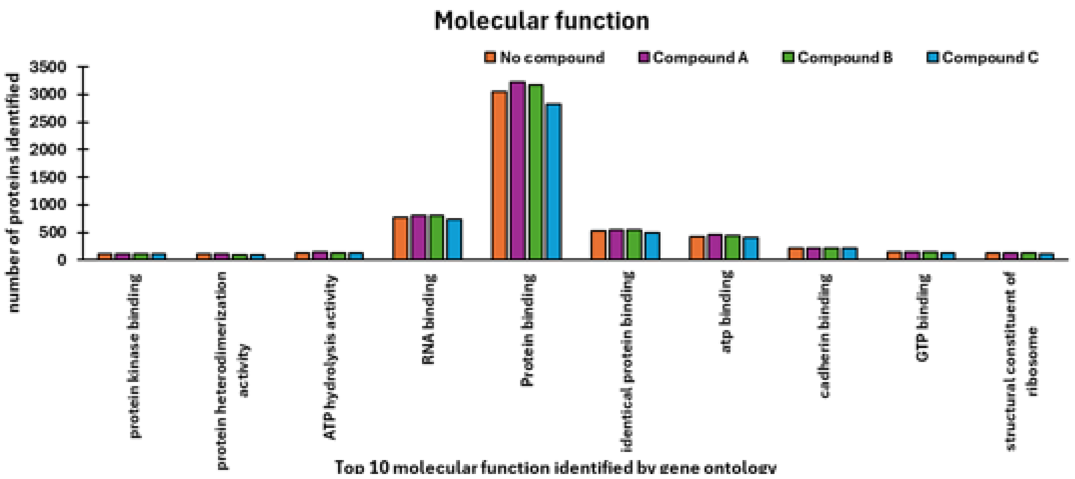


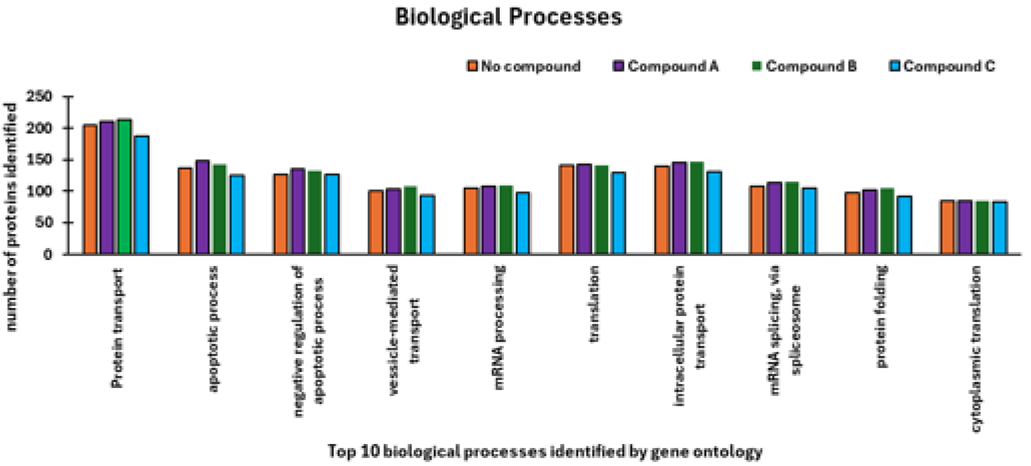


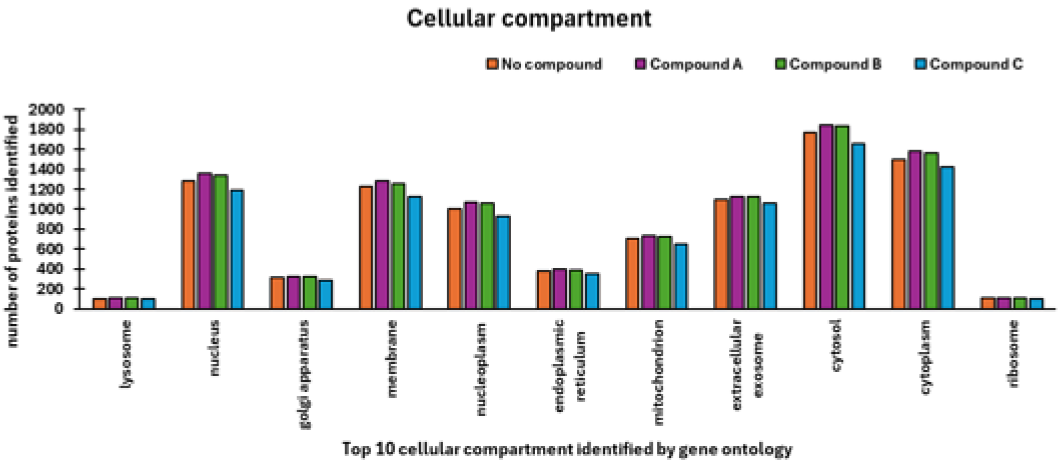


**Figure S3**. Gene ontology plots with respect to molecular function (top), biological processes (middle), and cellular components (bottom).

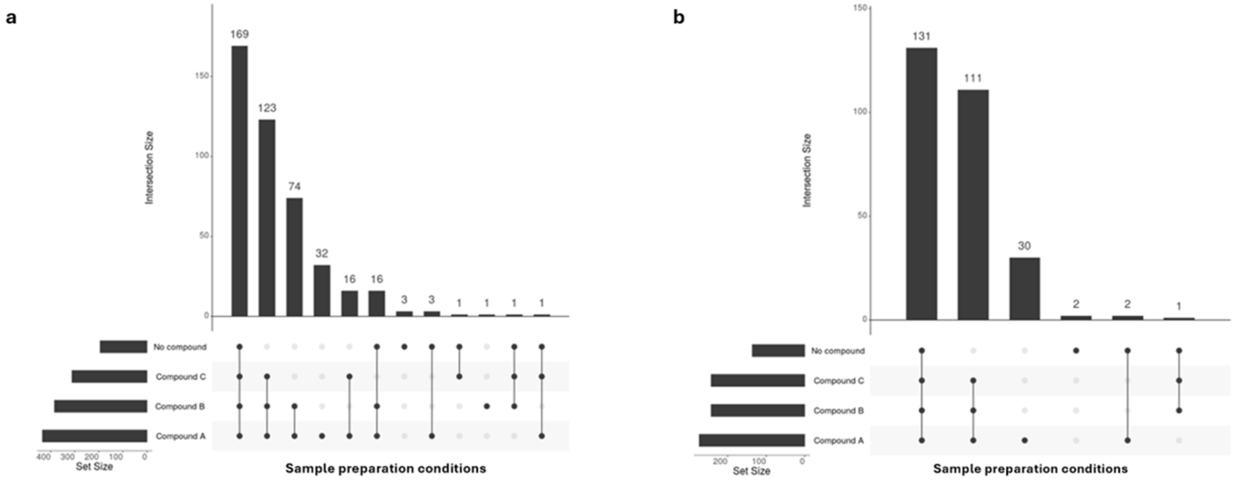


**Figure S4.** Upset plots describing overlaps in (a) cysteine-containing precursor identifications (**a**) and (**b**) proteins from the cysteine-containing precursors for all sample preparation conditions.

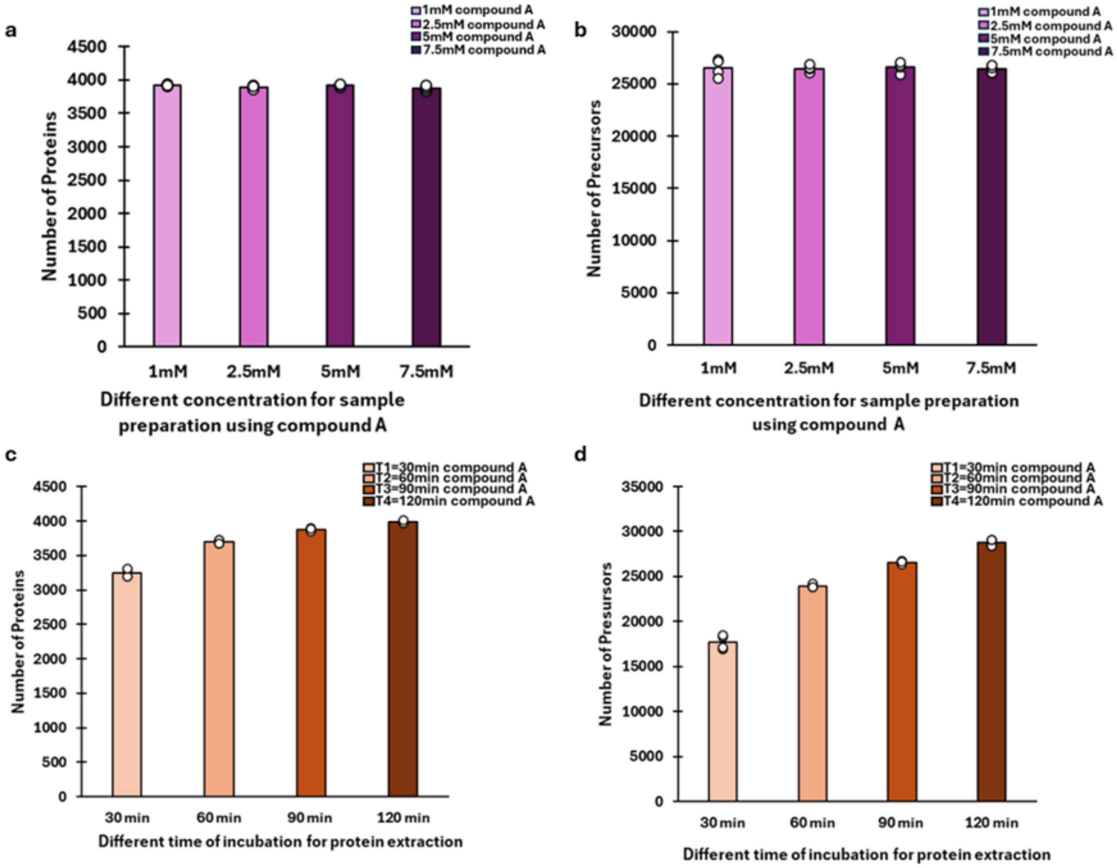


**Figure S5**. Compound A (3,4-diaminobenzoic acid) sample processing optimization. The number of (**a**) proteins and (**b**) precursors identified with various compound A concentrations during sample preparation. The number of (**c**) proteins and (**d**) number of precursors identified with different incubation times and 1mM compound A concentration. N = 4 for each condition.
